# Tau oligomer heterogeneity and associated protein profile in slowly versus rapidly progressive Alzheimer’s disease

**DOI:** 10.64898/2025.12.10.693456

**Authors:** Tayyaba Saleem, Wiebke Möbius, Matthias Schmitz, Angela da Silva Correia, Carolina Thomas, Sezgi Canaslan, Peter Hermann, Stefan Goebel, Saima Zafar, Elisabeth Root, Christine Stadelmann, Olivier Andreoletti, Michael Hoppert, Tiago Fleming Outeiro, Isidre Ferrer, Neelam Younas, Inga Zerr

## Abstract

Rapidly progressive Alzheimer’s disease (rpAD) is a rare but devastating clinical variant characterized by abrupt cognitive decline, yet the molecular features underlying this phenotype remain unknown. Tau oligomers (TauO) are key mediators of tau toxicity, but whether their biochemical properties differ across AD subtypes has not been examined in human brain. We isolated endogenous TauO from frontal cortex of well-characterized control, slowly progressive AD (spAD), and rpAD cases using T22 immunoprecipitation and performed ultrastructural, biochemical, and proteomic characterization. rpAD TauO displayed compact, densely aggregated morphology and exhibited the highest levels of disease-associated phosphorylation (pS396, pS422). Label-free proteomics revealed that control and spAD shared a robust TauO interactome enriched for translation, proteostasis, mitochondrial metabolism, and vesicle trafficking. Strikingly, these modules were absent in rpAD, which instead showed selective enrichment for aldehyde detoxification, amino-acid and carbon metabolism, and actin-regulatory pathways. rpAD TauO demonstrated increased association with SERPINA1, ALDH9A1, MAPRE3, DPYSL2/3, and NFASC, and reduced association with MRPL17 and C9. Functionally, rpAD TauO induced the strongest toxicity in SH-SY5Y cells. Together, these findings indicate that rpAD likely harbors a biochemically distinct TauO species, defining a molecular signature that may underpin its fulminant clinical progression and support the development of subtype-specific therapeutic strategies.

## 1. Introduction

Alzheimer’s disease (AD), the most common neurodegenerative disorder and the leading cause of dementia worldwide, is characterized by progressive cognitive impairment and extensive neuronal loss. The pathological hallmarks of AD are extracellular β-amyloid (Aβ) plaques and intracellular neurofibrillary tangles (NFTs) composed of hyperphosphorylated tau protein ^1–3^. In recent years, the accumulation of Aβ plaques and subsequent NFT growth have been shown to be responsible for downstream synaptic dysfunction and neuronal loss, indicating the onset and progression of AD-associated symptoms ^2,4^.

Since the postulation of the amyloid cascade hypothesis in the 1990s, the notion of direct Aβ-driven toxicity has dominated the field. However, the primary clinical and pathological findings have been questioned over the years. Notably, Aβ burden does not correlate well with disease severity or progression, and ∼30% of elderly individuals harbor substantial Aβ pathology without cognitive impairment ^5^. Moreover, numerous anti-Aβ therapeutics have failed in clinical trials, prompting renewed focus on tau pathology as a more proximal driver of neurodegeneration.

Under pathological conditions, tau, a microtubule-associated protein, becomes hyperphosphorylated and undergoes conformational changes that promote self-association into oligomers, paired helical filaments (PHFs), and NFTs ^6^.

Striking biochemical diversity exists in the forms of soluble, oligomeric and seed competent hyperphosphorylated tau in AD patients. In addition, some posttranslational modification sites (PTMs) are associated with the increased seeding activity of tau, leading to worse clinical outcomes ^7–9^. Among tau species, tau oligomers (TauO) are increasingly recognized as the most neurotoxic form, as they mediate synaptic dysfunction, axonal transport impairment, and neuron loss in both in vitro and in vivo models ^10^. TauO also exhibit prion-like properties, propagating between cells and promoting further aggregation of native tau ^11,12^.

The multiple tau biochemical states and strains may explain the clinical heterogeneity in AD phenotypes. Slowly progressive AD (spAD) and rapidly progressive AD (rpAD) differ not only in their clinical trajectory but also in their molecular features, including tau PTMs and conformational states ^5^. Emerging data suggest that tau conformers may form distinct strains with unique pathogenic properties, akin to prions ^13^. To test this hypothesis, we isolated TauO from human frontal cortex tissue of individuals belonging to clinically well-characterized groups, namely, spAD, rpAD, and age-matched controls. We examined their morphology via transmission electron microscopy (TEM), assessed tau phosphorylation patterns in frontal cortex lysates, and characterized their copurified proteome via quantitative mass spectrometry. Our goal was to perform a clinicopathological correlation and identify whether TauO in rpAD differs structurally or molecularly from that in spAD and control brains, potentially offering mechanistic insight into their more aggressive clinical course.

## 2. Methods

### 2.1 Ethics statement

Frontal cortex tissue samples were collected postmortem from individuals with spAD (n=5), rpAD (n=5) and from age-matched non-demented controls (n=5). Tissues were sourced from the Institute of Neuropathology Brain Bank (HUB-ICO-IDIBELL Biobank) and the Biobank of Hospital Clinic-IDIBAPS, Spain. Additional control samples were obtained from the Department of Neuropathology at the University Medical Center Göttingen, Germany. All procedures complied with national legislation and institutional protocols and were approved by the relevant ethics committees (Spain: HUB-ICO-IDIBELL Biobank, Hospital Clinic-IDIBAPS; Germany: University Medical Center Göttingen, protocols Nr. 1/11/93 and Nr. 9/6/08).

### 2.2 Patient cohorts and spAD/rpAD subtype characterization

Patient selection and neuropathological assessments followed previously established protocols^14^. All AD patients (spAD and rpAD) exhibited advanced neurofibrillary pathology (Braak stage > V) and were free of coexisting neurodegenerative conditions. Diagnosis was confirmed through standardized neuropathological evaluation of 25 brain regions, including the cerebral cortex, thalamus, diencephalon, cerebellum, and brainstem, as described previously ^15^. Histological techniques included hematoxylin and eosin staining, Klüver-Barrera staining, and immunohistochemistry for β-amyloid, phosphorylated tau, GFAP, alpha-synuclein, TDP-43, ubiquitin, p62, and microglial markers. All rpAD patients fulfilled contemporary diagnostic criteria ^16^. The full Braak and Thal staging details are provided in supplementary Table 1S.

### 2.3 Cohort and assay allocation

TauO were isolated from 15 patients (control n=5, spAD n=5, rpAD n=5). Western blot PTM assessment, TauO isolation, and TEM were performed on all 15 samples. For discovery proteomics, the eluate per case was finite, and material was allocated prospectively across assays; at the time of LC-MS acquisition, we therefore included the samples with sufficient remaining eluate, yielding control n=3, spAD n=5, and rpAD n=3 (rpAD1, rpAD2, and rpAD4; Table S1). All proteomics samples were prepared with equalized input and processed under identical buffers with matched IgG controls.

### 2.4 Tissue homogenization and protein extraction for TauO

Frozen frontal cortex tissues were homogenized at a 1:3 (w/v) ratio in ice-cold phosphate-buffered saline (PBS) supplemented with a protease inhibitor cocktail (1 tablet per 50 mL, Roche). Homogenization was performed via a mechanical tissue lyser (Qiagen) to ensure complete cellular disruption. The lysates were subsequently centrifuged at 9,279 × g for 10 minutes at 4 °C. The resulting supernatants (the PBS-soluble fraction) were collected and aliquoted for subsequent biochemical analyses.

### 2.5 Western blot analysis of TauO

Western blotting was performed to confirm the presence of high-molecular-weight tau oligomers in both total lysates and immunoprecipitated fractions. The samples were diluted in 4× Laemmli sample buffer (Bio-Rad) without boiling to preserve oligomeric structures. Proteins were separated via NuPAGE on 4-12% Bis-Tris precast gels (Invitrogen) in 1× MOPS SDS running buffer at 80 V for 10 minutes followed by 120 V for ∼1 hour. Proteins were transferred to nitrocellulose membranes (0.45 µm, GE Healthcare) via wet transfer (97 V, 1 hour, 4 °C). The membranes were blocked in 5% BSA and probed overnight at 4°C with primary T22 antibody (1:1000) followed by HRP-conjugated anti-mouse secondary antibody. Detection was performed via enhanced chemiluminescence (ECL) and imaging via a ChemiDoc imaging system (Bio-Rad).

### 2.6 Immunoprecipitation and isolation of TauO

Tau oligomers (TauO) were enriched from PBS-soluble brain lysates via immunoprecipitation using the tau oligomer-specific antibody T22 (Millipore). Tosyl-activated magnetic Dynabeads (Thermo Fisher Scientific) were conjugated with 20 µg of T22 antibody (1.0 mg/mL) in 0.1 M borate buffer (pH 9.5) by overnight incubation at 37 °C. The beads were washed in 0.2 M Tris buffer (pH 8.5) containing 0.1% BSA to remove unbound antibody and block nonspecific binding sites.

The PBS-soluble lysates were incubated with the antibody-conjugated beads under gentle rotation for 1 hour at room temperature. After incubation, the beads were washed 3× in PBS to eliminate unbound material. Bound TauO was eluted with 0.1 M glycine (pH 2.8), and the eluate was immediately neutralized with 1 M Tris-HCl (pH 8.0). The eluted fraction was then concentrated and buffer-exchanged via Microcon centrifugal filters (25 kDa MWCO, Millipore) at 14,000 × g for 25 minutes at 4 °C. Finally, the TauO protein was resuspended in PBS and quantified via a BCA protein assay (Thermo Fisher).

### 2.7 Transmission electron microscopy (TEM)

For negative staining of TauO for electron microscopy, 10 µL of the TauO sample mixture was applied to a paraffin film. A glow-discharged carbon-coated copper grid (400 mesh) was then placed onto the drop to allow adsorption of the sample. Subsequently, 10 µL of 0.25% glutaraldehyde was added to the drop and incubated for 1 min to fix the sample. The grid was washed briefly three times by dipping in PBS to remove excess fixative. Next, the grid was incubated for 30 seconds with 2% uranyl acetate stain. Importantly, no washing was performed after uranyl acetate incubation; instead, excess stain was carefully wicked off, and the grid was air-dried before further analysis. Morphological analysis of TauO was performed via transmission electron microscopy. The stained grids were examined to assess features such as size, shape, circularity, and the presence of electron-dense aggregates in TauO derived from control, spAD, and rpAD samples.

### 2.8 Cell culture and treatment

Cells were maintained in T75 flasks with DMEM supplemented with 10% FBS and 1% penicillin-streptomycin. At 70–90% confluency, cells were trypsinized, washed with 1× PBS, and seeded into 96-well plates at a density of 1 × 10⁴ cells/well. After incubation for 18-24 h at 37 °C, cells were treated with brain-derived tau oligomers (TauO) from experimental and control groups for 18hours.

### 2.9 MTS assay

Cell viability was assessed using the MTS assay (Abcam, ab197010) according to the manufacturer’s instructions. Briefly, culture medium was replaced with fresh medium prior to adding MTS reagent [3-(4,5-dimethylthiazol-2-yl)-5-(3-carboxymethoxyphenyl)-2-(4-sulfophenyl)-2H-tetrazolium, inner salt]. Cells were incubated for 1 h at 37 °C to allow the formation of a soluble formazan product by metabolically active cells. Absorbance was measured at 490 nm using a Perkin Elmer Wallac 1420 Victor microplate reader (GMI, USA). Background absorbance from control wells was subtracted from experimental readings to obtain final values.

### 2.10 Label-Free Quantification Mass Spectrometry (LFQ-MS) Analysis

TauO fractions were resolved by short-term SDS‒PAGE (4‒20% Bis‒Tris) and subjected to in-gel tryptic digestion. Peptide mixtures were spiked with Biognosys iRT standards and analyzed on a Thermo Exploris 480 Orbitrap using a 22-variable window DIA method (60 min gradient, 250 ng equivalent per run). Each biological replicate was analyzed in technical triplicate.

The data were processed in Spectronaut v19.0 (Biognosys) via the Pulsar search engine against the UniProtKB *H. sapiens* reference proteome (release 08/2023) supplemented with a 53-protein contaminant database. Protein and peptide identification was controlled at a 1% false discovery rate (FDR). Quantification was performed by DIA using up to 6 fragments per peptide and up to 10 peptides per protein, with dynamic retention time alignment, dynamic mass recalibration, and quartile normalization. Spectronaut’s built-in global data imputation was applied to the final results table. For downstream filtering, only proteins supported by ≥2 stripped peptides in the experiment-wide report were retained. To minimize nonspecific background, matched isotype IgG immunoprecipitations were performed for each group. Proteins enriched ≥2-fold in TauO (T22 IP) relative to their matched IgG controls (linear space; equivalent to log2FC ≥ 1) were considered copurified with TauO. LFQ intensities (arbitrary MS1-based units) were log2-transformed and z scored for heatmap visualization; for boxplots, data are displayed as log10 LFQ intensities (arbitrary units).

### 2.11 Humanized 3xTg mouse model of combined Aβ and tau pathology

The triple-transgenic (3xTg) mouse model, originally developed by Oddo et al., carries mutations in the APP, PS1, and tau genes, recapitulating key features of combined Aβ and tau pathology ^17^. In this study, 3- to 4-month-old 3xTg mice received stereotactic injections of 20 µL of 10% (w/v) cortical brain homogenate from an Alzheimer’s disease (AD) patient, which was targeted to the thalamus. The homogenate was prepared in PBS and clarified by centrifugation prior to inoculation. Both inoculated (AD) and noninoculated control mice were sacrificed at 3, 6, 9, and 12 months post-injection. After decapitation, the cortical tissues were rapidly collected and snap-frozen in liquid nitrogen for further analysis. A sample size of n = 4 per group was chosen on the basis of practical considerations and precedent from prior 3xTg studies.

### 2.12 Gene ontology (GO) term analysis

Functional enrichment and network analyses were performed in Metascape (Homo sapiens). Input lists were derived from tau-oligomer co-purified brain proteomes and included: (i) unique sets per condition (Control-only, spAD-only, rpAD-only, shared spAD-rpAD), and (ii) where indicated, core per-condition lists defined purely by identification confidence (proteins with ≥2 unique peptides within a condition, independent of abundance). Gene identifiers were supplied as HGNC symbols. For all analyses, the background was the union of all quantified proteins detected in the experiment, to control for mass-spectrometry detection bias. Enrichment was run against GO Biological Process, KEGG, Reactome, and CORUM with Metascape defaults (minimum overlap=3, p-value cutoff = 0.01; Benjamini-Hochberg FDR applied by Metascape). Redundant term clusters were consolidated using Metascape’s default hierarchical reduction. Protein-protein interaction (PPI) modules were identified with MCODE using Metascape defaults (node degree and score thresholds as provided; network size bounds 3-500) on the Physical Core interactome. Disease enrichment used DisGeNET within Metascape. Outputs included heatmaps of enriched terms and PPI networks colored by MCODE cluster; exported PDFs/CSVs were used for figure assembly and reporting. All analyses were run with the same organism setting (Homo sapiens) and the same custom background to ensure comparability across lists.

### 2.13 Statistical analysis

Statistical analysis for mass spectrometry data was performed using Python (3.11.9), and GraphPad Prism (version 9). Libraries used in the Python included pandas, matplotlib and seaborn. Group comparisons were performed in GraphPad Prism v9. For two-group comparisons, Welch’s t test (parametric) or the Mann-Whitney U test (nonparametric) was applied as appropriate. Statistical significance was defined as p<0.05 on the basis of log2-transformed LFQ intensities.

## 3. Results

Given the clinical and pathological heterogeneity between spAD and rpAD, we explored whether TauO isolated from these subtypes and control brains exhibit distinct structural and molecular characteristics. To explore the molecular differences in tau oligomers across AD subtypes, we conducted a multistep analysis integrating biochemical, ultrastructural, and proteomic approaches.

### 3.1 Distinct patterns of Tau oligomers in AD subtypes

Western blotting of PBS-soluble brain lysates with the oligomer-selective antibody T22 revealed high-molecular-weight (HMW) tau species across groups, with variable intensity in the control group and more prominent ∼150 kDa bands in the spAD group (Supplementary Fig. 1S). rpAD showed a distinct but overall less intense HMW pattern (Supplementary Fig. 1S). Following T22 immunocapture, HMW TauO was confirmed in eluates from the control, spAD, and rpAD groups (***Figure 1***), indicating successful enrichment.

**Figure 1.**
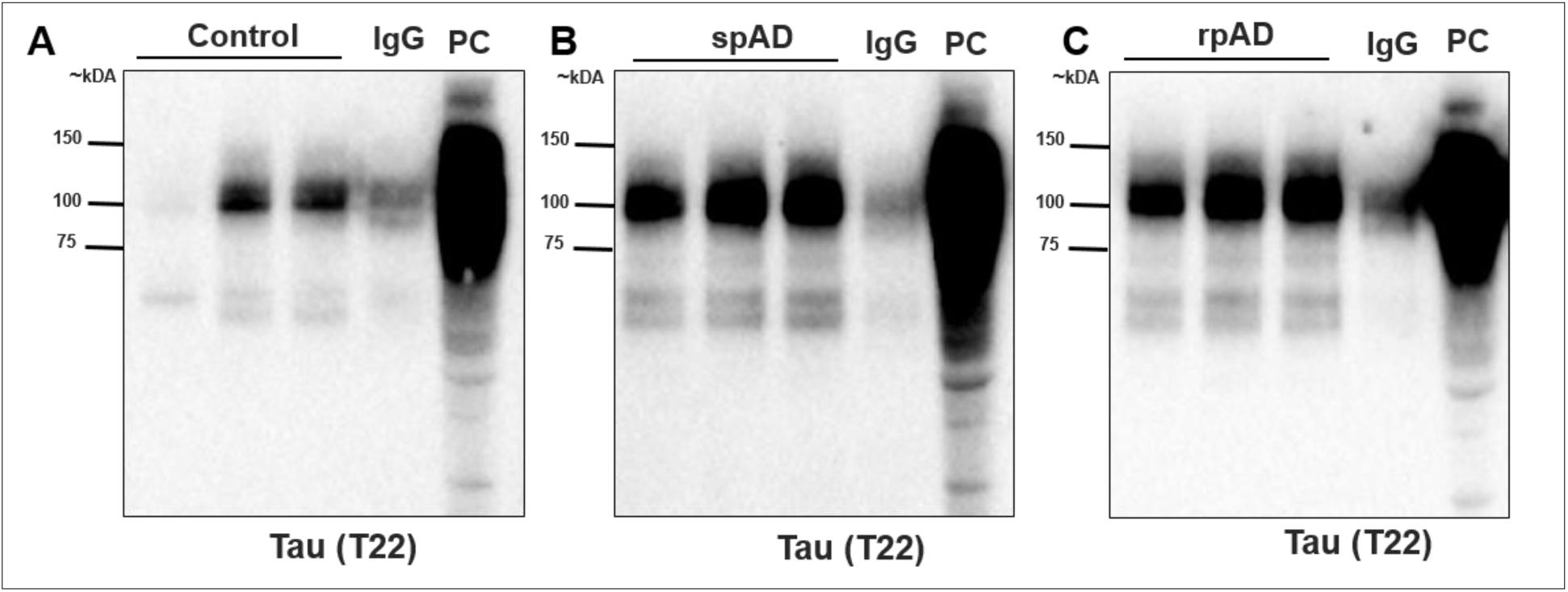
Western blot analysis of isolated TauO in control, spAD, and rpAD brain samples (frontal cortex) via the T22 antibody. TauO were immunoprecipitated from human brain homogenates and probed with a T22 antibody. HMW TauO particles were successfully isolated from all three groups. Representative immunoblots showing tau oligomer profiles in control (A), spAD (B), and rpAD (C) samples. IgG lanes represent isotype IgG negative controls immunoprecipitation, and PC lanes correspond to the positive control. Representative blots from 3 of 5 analyzed samples (n = 15 (5 control, 5 spAD, 5 rpAD)).

### 3.2 Tau oligomer morphology differs across disease subtypes

Negative-stain TEM of T22-enriched soluble fractions revealed clear qualitative differences in oligomer appearance across groups (*Figure 2*). Control samples showed the lowest oligomer abundance. Many fields contained no visible particles, and representative fields displayed predominantly small, uniformly spherical oligomers with only rare hollow-centered (annular) structures and minimal aggregation. spAD samples exhibited the greatest morphological diversity, including small spherical particles, more frequent annular oligomers, irregular electron-dense aggregates, and occasional membrane-like or vesicular structures. rpAD samples also contained spherical, annular, and aggregated particles; however, their fields were characterized by high oligomer abundance and dense particle clustering, with a visually more uniform population of small round oligomers and fewer distinct structural subclasses compared with spAD. The bottom row shows zoomed-in regions highlighting representative particle morphology in each group.

**Figure 2.**
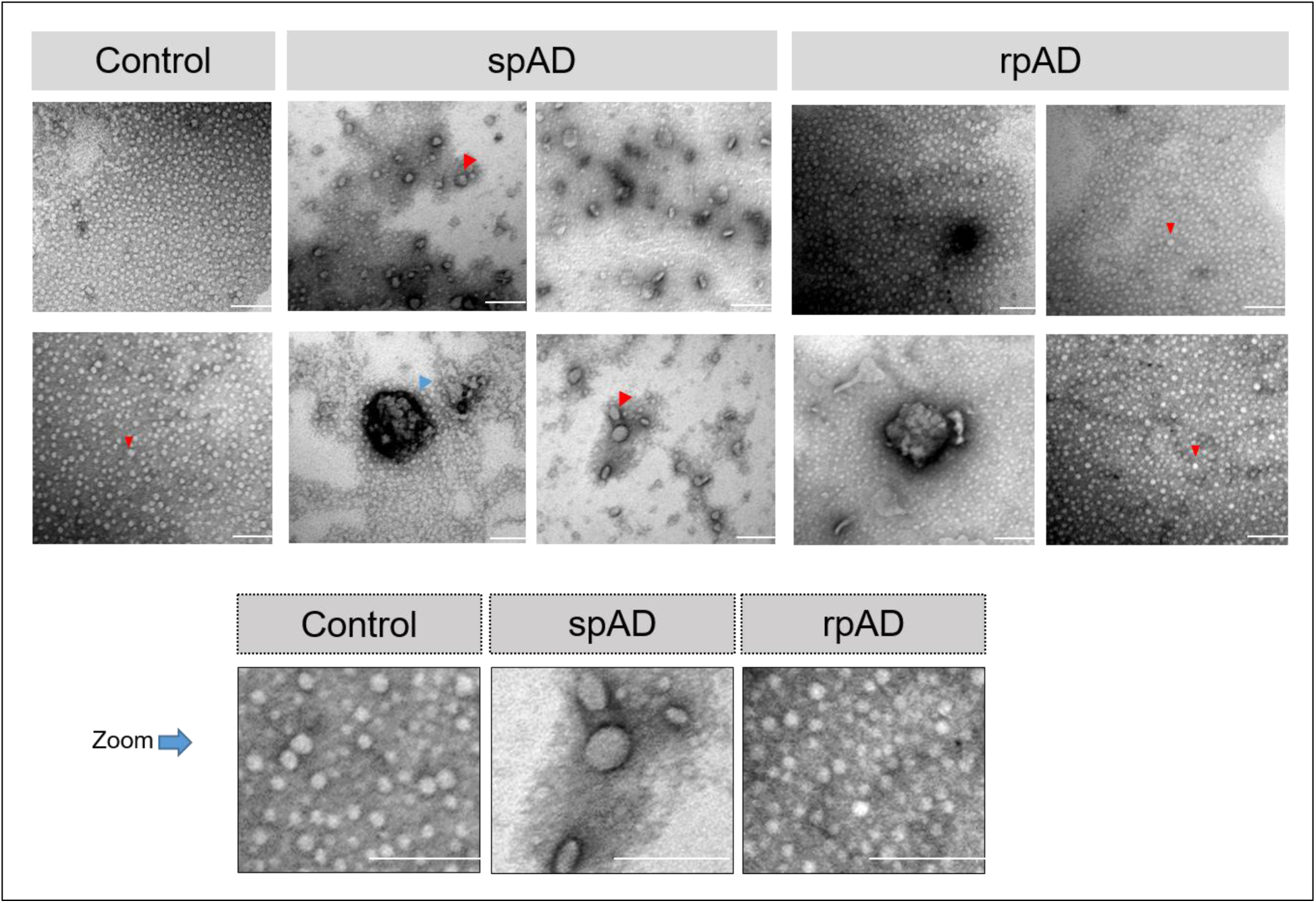
Negative-stain TEM of endogenous T22-enriched tau oligomers from control, spAD, and rpAD frontal cortex. Representative micrographs from T22-enriched soluble fractions (n = 5 per group). Control samples show predominantly small spherical particles with infrequent annular structures (red arrowheads). spAD samples display a broader range of particle morphologies, including spherical, annular (red arrowheads), and irregular aggregated forms, as well as occasional membrane-like material (blue arrowhead). rpAD samples show abundant, densely packed small spherical oligomers with additional annular and aggregated species. Bottom panels show zoomed-in regions from representative images. Scale bars = 100 nm.

### 3.3 Evaluation of tau post-translational modifications (PTMs) in AD subtypes

Tau oligomerization is strongly influenced by PTMs, particularly phosphorylation, which disrupts microtubule binding and exposes aggregation-prone domains. To assess potential subtype-specific differences, we screened multiple tau phosphorylation sites in urea/thiourea lysates. A schematic of the tau domains and epitopes examined is shown in Figure 3A. Across the sites analyzed (Tau-5, S198, S199, T205, T231, and S404), no significant group differences were detected (Supplementary Fig. 2S). In contrast, phosphorylation at S396 and S422 displayed the most marked alterations (Figure 3 B-C). The level of phosphorylation at S396 did not differ significantly between spAD patients and controls, but rpAD was markedly greater than both spAD patients (p = 0.0021) and controls (p = 0.0004). Similarly, phosphorylation at S422 was not significantly altered in spAD versus controls, yet rpAD exhibited significantly elevated levels compared with both groups (p = 0.0327 vs spAD; p = 0.0125 vs control). Controls exhibited minimal phosphorylation at these epitopes, which was consistent with physiological tau regulation. Hyperphosphorylation at these sites in AD may destabilize microtubule binding and facilitate tau oligomer formation, in line with previous mechanistic studies ^18^.

**Figure 3.**
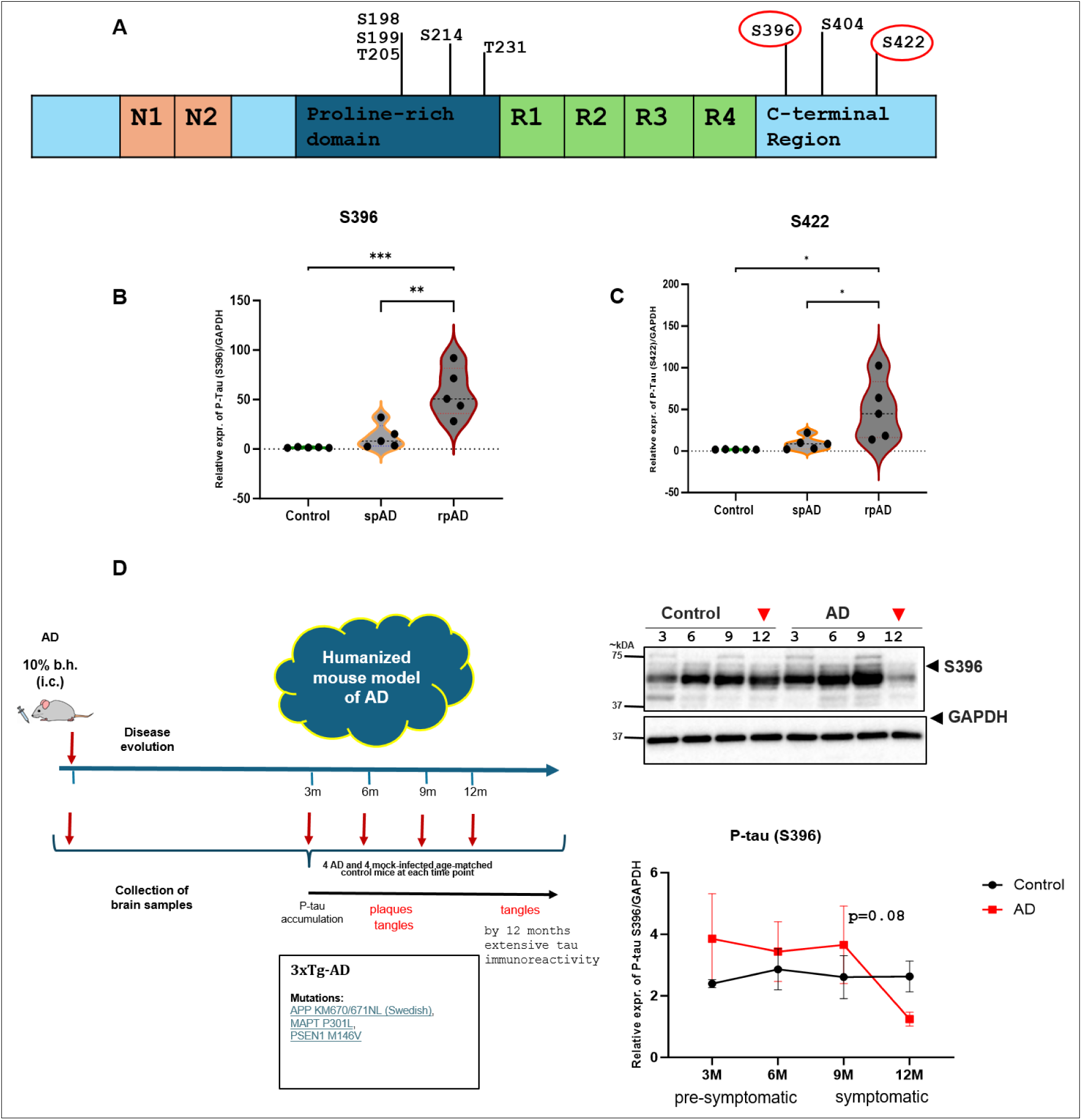
Tau PTMs in control, spAD, and rpAD strains and validation in a 3xTg mouse model. (A) Schematic of tau protein domains indicating the phosphorylation sites examined; sites showing significant differences are highlighted in red circles. (B-C) Violin plots of the pS396 and pS422 intensities in the control (n = 5), spAD (n = 5), and rpAD (n = 5) groups. (D) Western blot validation of pS396 expression in cortical tissue from 3xTg AD mice inoculated with AD brain homogenate (n = 4 per time point) compared with non-inoculated controls (n = 4). GAPDH served as a loading control. One-way ANOVA was used for human groups (B-C), and unpaired t tests with multiple-comparison correction were applied at each time point in mice (D). *p < 0.05, **p < 0.01, ***p < 0.001.

### 3.4 Age-dependent changes in S396 phosphorylation in 3xTg mice

To investigate the temporal regulation of pS396, we examined cortical lysates from 3xTg AD mice inoculated with AD brain homogenates (Figure 3D). Immunoblotting revealed the progressive accumulation of pS396 tau at early time points (3-9 months post-inoculation), which was consistent with the development of tau pathology. By 12 months, the pS396 levels had declined relative to earlier peaks. This reduction may reflect sequestration of hyperphosphorylated tau into insoluble aggregates or loss of tau-expressing neurons with disease progression ^19^. Similar late-stage declines in soluble phosphorylated tau have been described in other AD models, supporting the concept of a dynamic shift in tau solubility during disease evolution ^20^.

### 3.5 Brain-derived TauO induces toxicity in a neuronal cell model

To evaluate the bioactivity of brain-derived TauO, SH-SY5Y neuroblastoma cells were treated with recombinant tau pre-incubated with TauO isolated from control, spAD, and rpAD brain homogenates. Successful seeding and aggregation of recombinant tau were verified by T22 immunoreactivity, confirming that all preparations retained oligomeric competence. In contrast to recombinant tau monomers did not affect cell viability, exposure to 0.25 μM TauO from both spAD and rpAD samples caused a marked reduction in the MTS signal (***Figure 4***). The decrease in metabolic activity was more pronounced for rpAD TauO, showing a trend toward greater toxicity compared with spAD TauO, although this difference did not reach statistical significance (p=0.27). This graded pattern from no effect in monomer treated to clear toxicity in spAD and the strongest response in rpAD suggests a hierarchy of bioactivity among human brain-derived TauO species. These findings align with previous reports describing enhanced neurotoxicity and seeding capacity of soluble tau assemblies derived from AD brains ^21,22^.

**Figure 4.**
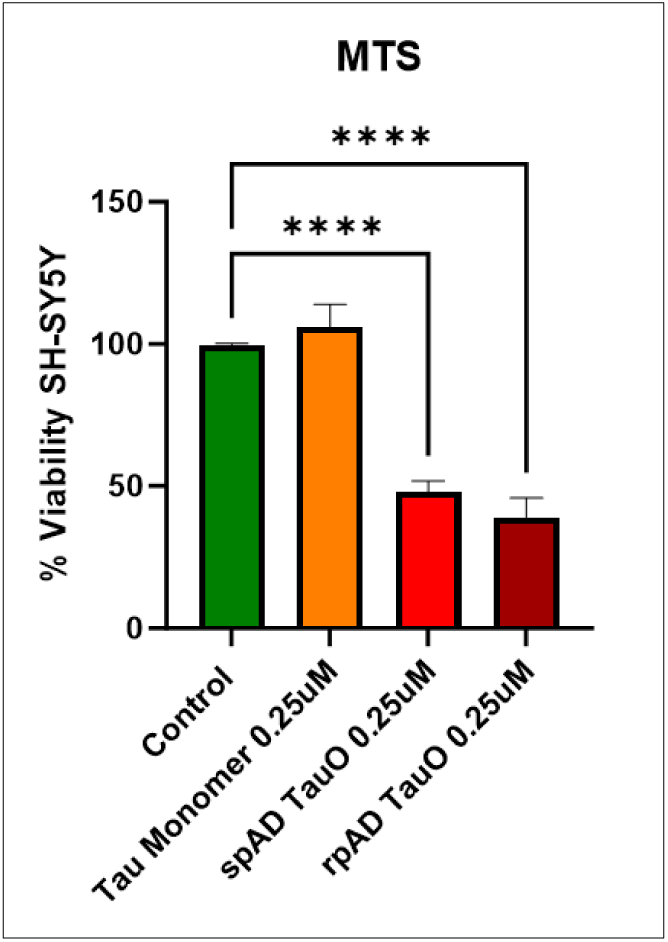
Cytotoxic effects of brain-derived TauO in SH-SY5Y cells. MTS assay quantification of cell metabolic activity following exposure to 0.25 μM recombinant tau monomers (orange) or brain-derived TauO (red = spAD; dark red = rpAD). Control (green) show maximal viability. TauO treatment significantly reduced the MTS signal relative to that of the controls (p < 0.0001 for spAD; p < 0.0001 for rpAD). The viability of the rpAD TauO group tended to be lower than that of the spAD TauO group (p = 0.27). The bars represent the means ± SEMs.

### 3.6 Comparative mapping of proteins co-purified with TauO

To characterize the subtype-specific TauO proteome, we used label-free quantitative mass spectrometry (Figure *5*a). 2,388 proteins were identified across all the groups. Venn diagram analysis revealed 556 proteins shared among the control, spAD, and rpAD groups, likely representing a copurified proteome (Figure *5*b). Remarkably, the rpAD samples presented the greatest number of uniquely enriched proteins (n = 1101), indicating a distinct composition of TauO-associated proteins. The spAD and control samples contained 81 and 198 unique proteins, respectively.

**Figure 5.**
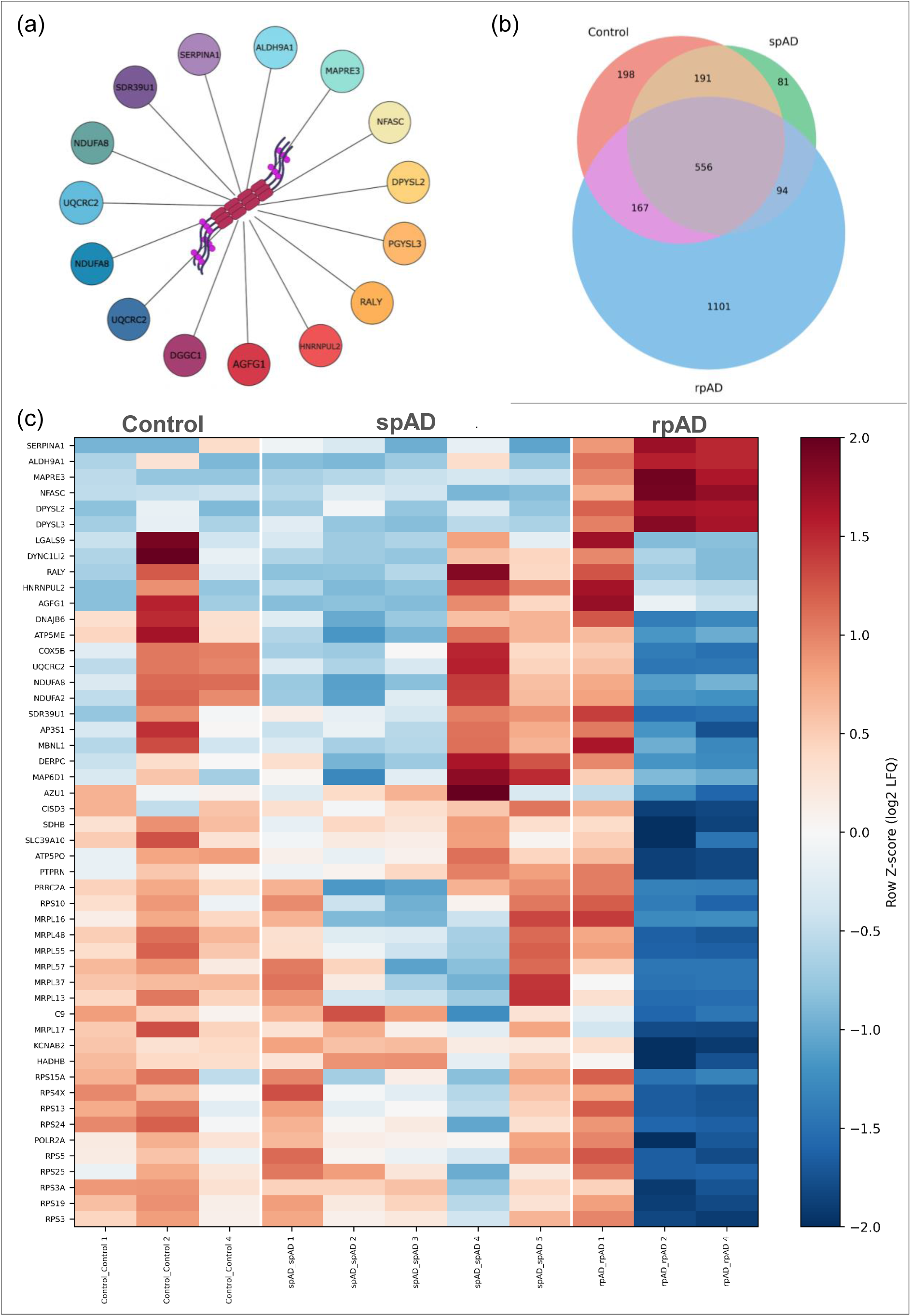
Core tau-oligomer (TauO)-associated proteins identified across disease groups. (A) Representative schematic showing selected proteins detected in TauO-immunoprecipitated fractions. Proteins are depicted as colored nodes surrounding TauO (center) to illustrate the diversity of co-purifying interactors. (B) Venn diagram summarizing overlap among proteins retained under uniform inclusion criteria (≥2 peptides; ≥2-fold enrichment versus IgG control). Counts: Control-only =198; spAD-only = 81; rpAD-only = 1,101; shared across all = 556; pairwise overlaps as indicated. (C) Heatmap of log₂-transformed, row Z-scored LFQ intensities for the core set of TauO-associated proteins across Control, spAD, and rpAD samples. Each column represents an individual case; rows correspond to quantified proteins. The color scale reflects relative abundance within each protein (n = 11; Control = 3, spAD = 5, rpAD = 3). Together, the panels depict group-specific variation in TauO-associated proteomes, highlighting extensive expansion of unique interactors in rpAD alongside quantitative divergence in shared core components.

A heatmap further illustrated clear groupwise clustering of the differentially enriched proteins that presented unique protein signatures in rpAD patients (***Figure 5***c). Despite the heterogeneity within the patient groups, the rpAD TauO samples presented distinct proteomic profiles, with notable enrichment of proteins such as SERPINA1, ALDH9A1, MAPRE3, NFASC, DPYSL2 and DPYSL3, and depletion of proteins such as MRPL17, and C9.

### 3.7 Selective loss of mitochondrial and translational signatures within the core TauO-associated proteome in rpAD

Metascape analysis of the core TauO-associated proteomes revealed that although the three groups share a substantial fraction of interacting proteins, the composition of enriched biological pathways differed sharply across conditions. The Circos plot showed that while a conserved core is present, the rpAD group contributed the largest condition-specific sector, suggesting divergence in pathway-level organization despite partial overlap at the protein level (Figure 6a). The enrichment heatmap demonstrated a striking functional separation between groups (Figure 6b). Control and spAD exhibited significant enrichment across the same set of thirteen pathways, including mitochondrial and metabolic functions (fatty acid degradation, lipid oxidation, respiratory chain complex I, carbon metabolism), vesicular trafficking (ER-to-Golgi and intra-Golgi transport, vesicle budding from the membrane), proteostasis-related modules (PA700 complex, ribonucleoprotein complex biogenesis, translation, 55S mitochondrial ribosome, PDCL/TRiC-CCT cooperation), and additional categories such as the NABA core matrisome etc. In contrast, rpAD showed no enrichment for any of these pathways and instead presented a completely distinct functional profile. Instead, the rpAD core interactome showed enrichment exclusively for five metabolic pathways absent from both control and spAD: aldehyde metabolic process, cysteine and methionine metabolism, protein homooligomerization, carbon metabolism, and L-amino-acid metabolic process. No pathway was shared across all three groups, and neither control nor spAD displayed any uniquely enriched pathway. Network-based visualization of these enriched terms further resolved them into coherent functional modules (Figure 6c–d). Control and spAD clustered into dense modules corresponding to translation/ribosome/proteostasis, vesicle transport, and mitochondrial metabolism. The rpAD-specific pathways formed a small but clearly separated metabolic module. Coloring by -log10(P) highlighted that the shared modules carried the highest statistical support.

**Figure 6:**
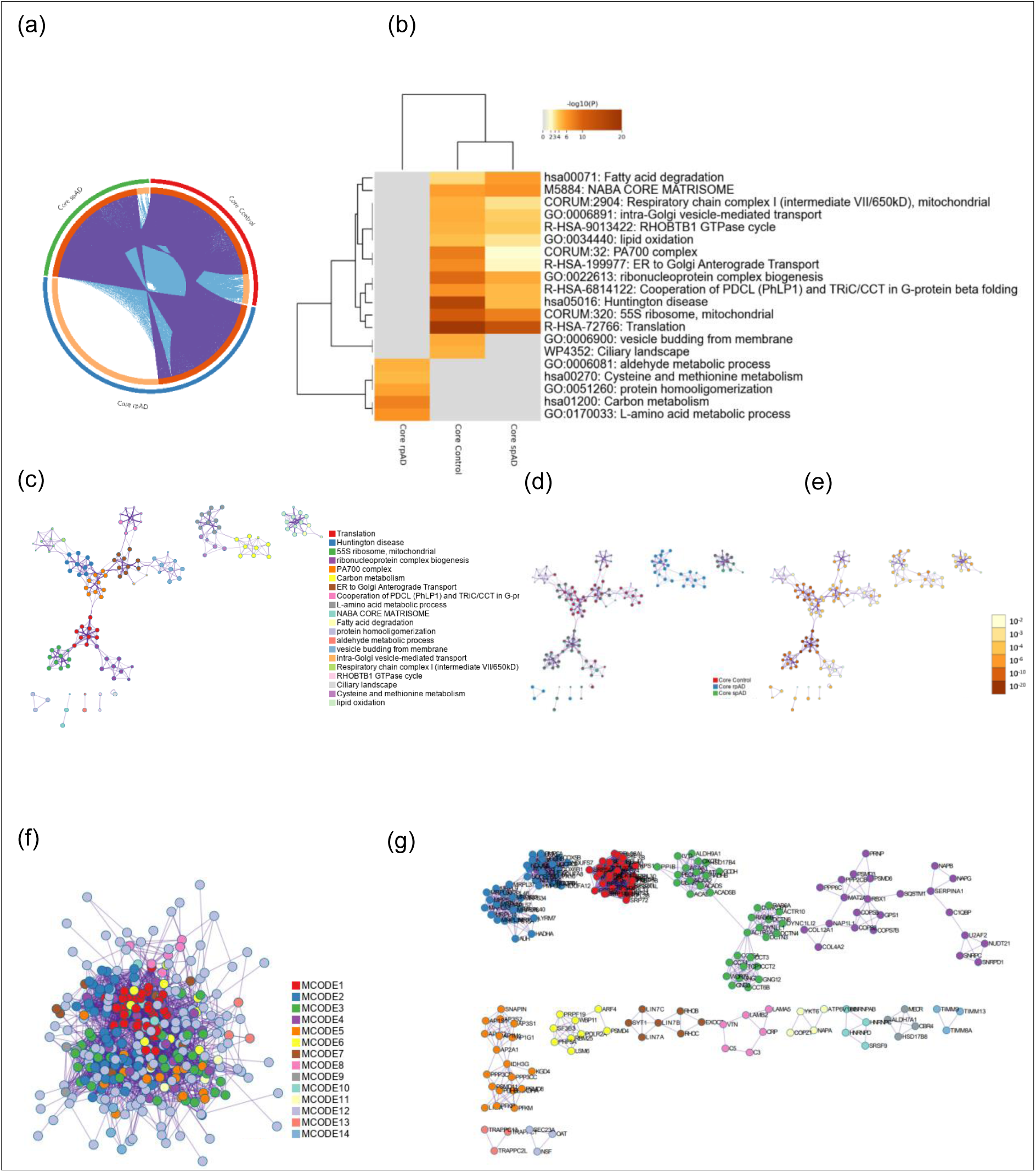
Metascape analysis of the core TauO-associated proteome across control, spAD, and rpAD groups. (a) Circos plot showing the overlap between the core TauO-associated proteins identified in control, spAD, and rpAD groups. Each outer ring represents one condition, and ribbons indicate shared proteins. A large rpAD-specific sector indicates condition-dependent divergence within the core interactome. (b) Heatmap of enriched GO/Reactome pathways (-log₁₀P) derived from the core TauO-associated protein sets. control and spAD show enrichment across mitochondrial, proteostasis, vesicle-trafficking, and matrisome-related pathways, whereas rpAD displays enrichment restricted to a small set of metabolic processes. (c) Enrichment network in which each node represents an enriched GO/Reactome term. Node size is proportional to the number of input genes annotated to the term, and node color denotes its Metascape-defined cluster identity. Terms with similarity >0.3 are connected by edges, with thicker edges indicating stronger similarity. (d) The same enrichment network as in (c), now color-coded by statistical significance (-log₁₀P). Darker colors denote more significant enrichment. Highly significant clusters correspond to translation, ribosome organization, vesicle trafficking, and mitochondrial pathways enriched in control and spAD. (e) Protein-protein interaction (PPI) network of all core TauO-associated proteins, with node coloration indicating their group of origin (control, spAD, rpAD). The network reflects the shared vs. condition-specific contributions to each connected module. (f) MCODE-clustered PPI network, in which densely connected neighborhoods are assigned distinct colors representing functionally coherent protein modules. (g) Representative MCODE subnetworks, showing modules enriched for translation/ribosome assembly, mitochondrial and metabolic processes, and vesicle-trafficking components.

Protein-protein interaction analysis with MCODE revealed densely connected subnetworks that mirrored these pathway themes, with large clusters representing translation machinery, mitochondrial proteins, and trafficking complexes in control/spAD, and a smaller, distinct metabolic module in rpAD (Figure 6e-g). Together, these data demonstrate that the core TauO interactome is structurally shared but functionally divergent, with control and spAD maintaining robust enrichment across proteostasis, mitochondrial, and trafficking modules, whereas rpAD exhibits a complete loss of these signatures and instead shows selective enrichment in a small set of metabolic pathways.

### 3.8 Unique TauO interactome exhibit distinct functional signatures across groups

To define condition-specific TauO interactomes, we analyzed proteins uniquely co-purified with TauO in each group. The Circos plot showed minimal overlap between unique protein sets, indicating that each condition contains a largely distinct complement of TauO-associated proteins (Figure 7a). Functional enrichment analysis revealed striking differences in pathway composition across groups (Figure 7b). The rpAD-unique interactome showed the broadest functional representation, with significant enrichment across multiple metabolic categories, including dicarboxylic acid metabolism, carbohydrate metabolism, carbon metabolism, metabolism of nucleotides, nucleotide-sugar and deoxyribonucleotide metabolism, together with protein depolymerization and regulation of the actin cytoskeleton by Rho GTPases (SIG regulation).

**Figure 7.**
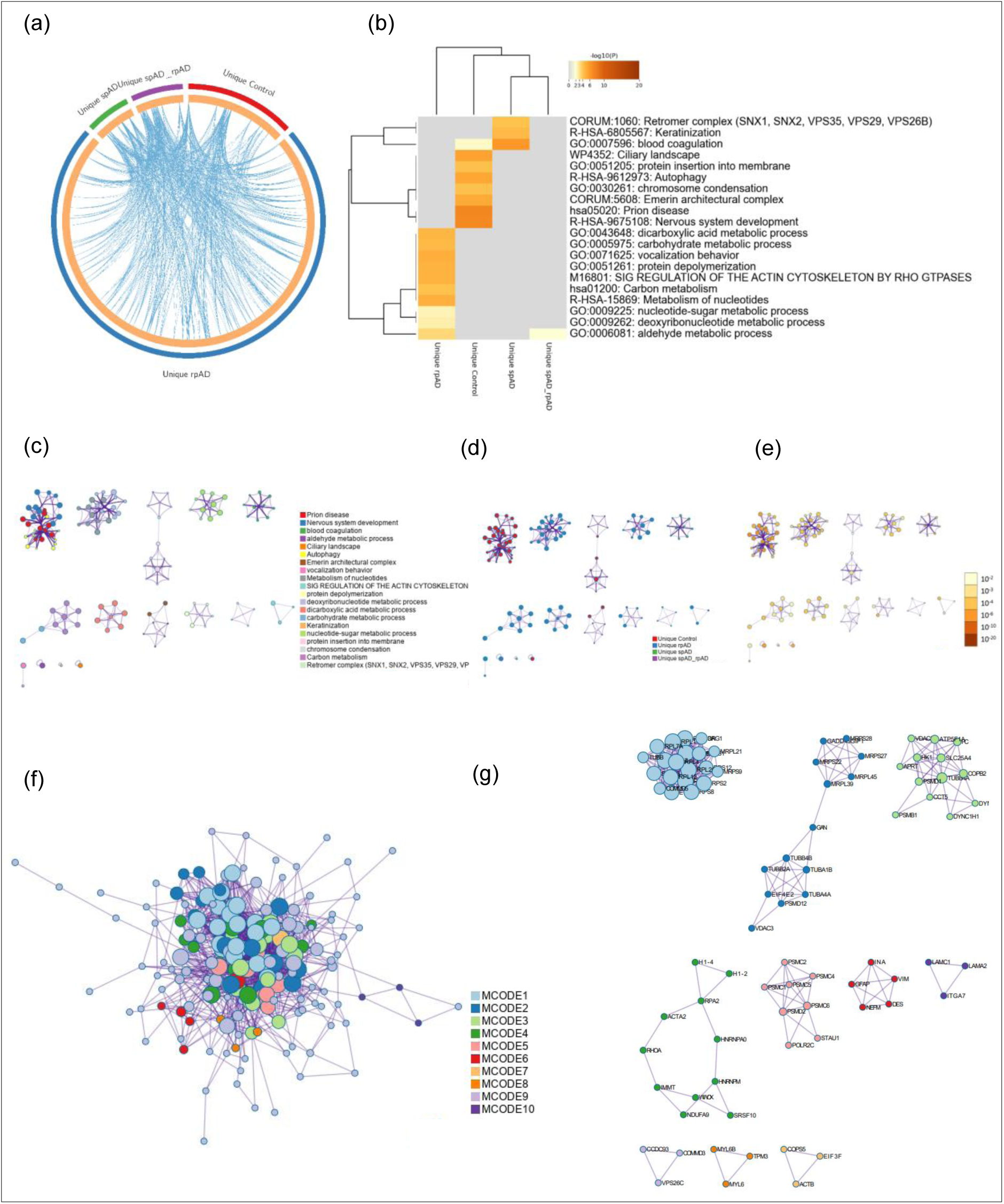
Pathway enrichment and network organization of unique TauO-interacting proteomes. (a) Circos plot showing the overlap between proteins uniquely identified in the control, spAD, and rpAD TauO-immunoprecipitated fractions. Each colored ribbon represents a group, and intersections indicate shared proteins. (b) Heatmap of enriched GO/Reactome pathways (-log₁₀P) derived from proteins unique to each condition. (c) Enrichment network in which each node represents an enriched term. Node size is proportional to the number of input genes annotated to that term, and node color denotes cluster identity. Terms with a similarity score >0.3 are connected by edges, with thicker edges indicating higher similarity. (d) The same enrichment network as in (c), but with nodes color-coded by statistical significance (-log₁₀P). Darker colors indicate more significant enrichment. (e) Protein-protein interaction (PPI) network constructed from all PPI relationships among genes unique to each group (f) MCODE-clustered PPI network generated from the control-unique protein set, showing densely connected subnetworks. (g) Representative MCODE subnetworks from the control-unique interactome, illustrating modules linked to autophagy, membrane insertion, nuclear architecture, and related processes.

The control-unique interactome exhibited enrichment for processes related to autophagy, protein insertion into membranes, chromosome condensation, ciliary functions, the Emerin architectural complex, and terms associated with prion disease and nervous system development. In contrast, the spAD-unique interactome showed a much narrower profile, with enrichment limited to the retromer complex and keratinization. Only a single pathway blood coagulation was shared between control and spAD but absent in rpAD, while aldehyde metabolic process represented the only pathway shared between rpAD and spAD.

Network-based pathway visualizations using Metascape further resolved these enrichments into discrete functional modules (Figure 7c-g). The rpAD-unique terms formed the largest and most densely connected clusters, dominated by metabolic and actin-regulatory modules. control-unique terms grouped into structural, nuclear, and cilia-associated modules, while the spAD-unique set formed a compact retromer-associated trafficking module. MCODE subnetwork analysis further resolved these enrichment profiles. In the control-unique interactome, major modules reflected translation and proteostasis (MCODE1), mitochondrial translation (MCODE2), chromosome and DNA organization (MCODE3-4), intermediate filament architecture, and ECM/laminin-integrin structural clusters. The rpAD-unique interactome yielded a single metabolically driven module centered on carboxylic acid and nucleotide metabolism, whereas spAD formed a compact retromer/retrograde trafficking module together with a coagulation cluster and a smaller mRNA-processing subnetwork. The shared spAD-rpAD subset produced a single amino-acid catabolic module. These MCODE-defined clusters further support and refine the pathway-level distinctions observed across conditions.

## 4. Discussion

Alzheimer’s disease is a clinically heterogeneous disorder, with distinct subtypes, such as spAD and rpAD, that differ in clinical course, neuropathological features, and molecular signatures ^23^. While spAD progresses over several years, rpAD is characterized by an abrupt cognitive decline and markedly shorter survival ^24^. The molecular determinants underlying this divergence remain poorly understood.

Here, we examined whether structural, biochemical, or proteomic features of TauO a key pathogenic intermediate differ between AD subtypes. Using immunoprecipitation, TEM, Western blotting, and label-free proteomics, we identified multiple convergent features that differentiate rpAD TauO from spAD and control TauO. Our results demonstrate that tau PTMs, particularly phosphorylation at S396 and S422, differ markedly between disease subtypes. While total tau levels were reduced in rpAD, possibly due to sequestration into insoluble aggregates, phosphorylation at S396 and S422 was significantly elevated in both spAD and rpAD, with rpAD showing the highest levels. Phosphorylation at S396 disrupts microtubule binding and promotes tau oligomerization and aggregation, whereas phosphorylation at S422 inhibits caspase-3 cleavage, potentially protecting oligomeric tau from degradation and facilitating its accumulation ^25,26^ Notably, both sites are associated with increased seeding activity and pathological progression in AD models. These PTMs may increase the formation and persistence of toxic oligomeric tau species, ^7,27^ particularly rpAD. The absence of these modifications in control TauO, despite detectable oligomeric bands by T22 and Western blot, suggests that phosphorylation rather than mere oligomerization is a key determinant of pathogenicity. In the 3xTg mouse model, we observed a temporal increase and subsequent decrease in pS396 tau levels, reflecting dynamic changes in tau pathology. The late-stage reduction likely results from sequestration into insoluble aggregates or neuronal loss, as reported in other AD models ^17,20^. These findings support the evolving nature of tau species during disease progression.

TEM analysis revealed that TauO morphology varied by diagnostic group. Control-derived TauO proteins are small and dispersed, which is consistent with physiological tau assemblies that stabilize microtubules ^28^. In support of this, a Drosophila tauopathy model showed that although hyperphosphorylated tau impaired synaptic and cytoskeletal function, granular tau did not correlate with increased toxicity. Furthermore, SH-SY5Y cells treated with control-derived TauO presented no significant decrease in viability, indicating that these oligomers are likely nontoxic and may play a stabilizing role in aging neurons ^28^. This aligns with prior work showing that brain-derived tau oligomers can share overall size and shape yet differ dramatically in seeding and toxic properties depending on conformation and PTM pattern ^29^. In contrast, spAD samples contained irregular, vesicle-like structures, suggesting early aggregation events ^6,30^. Interestingly, rpAD-derived TauO lacked these vesicle-like assemblies and instead exhibited small, dense oligomers. This may reflect a distinct aggregation trajectory involving rapid oligomer formation without progression to fibrils. Prior studies have shown that specific tau conformers can form structurally distinct “strains” with variable toxicity and propagation potential ^13^. Importantly, toxicity appears to be driven not simply by abundance but also by the conformational state and PTM profile, with hyperphosphorylation as seen in S422 and S396 ^31^. Our data therefore argue that morphology alone is insufficient to define pathological TauO; instead, a combination of PTMs and protein interactors likely tunes oligomer stability, cellular targeting and toxicity.

By integrating label-free proteomics with Metascape analysis, we delineated a shared “core” TauO interactome as well as condition-specific unique interactomes. Because the T22-IP captures co-purifying rather than strictly direct interactors, altered association may reflect pathway-level compartmentalization rather than binary binding. The core interactome was dominated in controls and spAD by modules related to translation (including cytosolic and mitochondrial ribosomes), proteostasis (PA700/19S proteasome complex, PDCL/TRiC-CCT chaperone machinery), mitochondrial metabolism and respiratory chain complex I, and vesicle trafficking (ER-to-Golgi and intra-Golgi transport, vesicle budding). These categories are consistent with earlier tau interactome studies showing strong enrichment of microtubule cytoskeleton, translational machinery and lysosomal components in brain-derived tau oligomer (BDTO) preparations and in phospho-tau interactomes ^32^.

In striking contrast, the rpAD core interactome showed complete loss of enrichment for these translation, proteostasis and vesicle-trafficking signatures. Instead, rpAD core TauO showed enrichment restricted to a small set of metabolic pathways aldehyde metabolism, cysteine and methionine metabolism, L-amino-acid and carbon metabolism and protein homooligomerization. The absence of any pathway that is jointly enriched across all three groups indicates that, despite considerable overlap at the individual protein level, the functional organization of TauO interactomes diverges sharply in rpAD. One interpretation is that in typical spAD, TauO remains predominantly embedded in canonical translation/mitochondrial/trafficking networks that may reflect both physiological tau functions and early stress responses, whereas in rpAD these networks are disengaged or overwhelmed, and TauO becomes selectively coupled to metabolic stress modules ^33,34^.

Analysis of proteins uniquely co-purifying with TauO in each group further sharpened these differences. The control-unique interactome was enriched for autophagy, protein insertion into membranes, nuclear and chromosomal organization, ciliary landscape, the Emerin architectural complex and pathways annotated to nervous system development and prion disease. The presence of autophagy and membrane insertion related modules is consistent with a physiological role for tau in vesicle trafficking and with cellular mechanisms that recognize and clear low levels of misfolded tau species ^31^.

The spAD-unique interactome was remarkably focused, with enrichment concentrated in the retromer complex and its associated retrograde transport pathways, together with keratinization and blood coagulation terms. Retromer dysfunction and endosomal trafficking defects are increasingly recognized as early features of AD and as contributors to both amyloid and tau pathology ^35^. The strong retromer signature in spAD-unique TauO suggests that oligomers in typical AD may preferentially engage endosomal/retrograde transport machinery, potentially facilitating propagation along defined neuronal pathways. By contrast, rpAD-unique TauO exhibited the broadest functional representation, dominated by metabolic pathways (dicarboxylic acid, carbohydrate and carbon metabolism; metabolism of nucleotides, nucleotide-sugars and deoxyribonucleotides; aldehyde metabolism) and regulation of the actin cytoskeleton by Rho GTPases, together with protein depolymerization. Enrichment of aldehyde metabolism and multiple carbon/energy pathways is notable given accumulating evidence that toxic aldehydes and impaired aldehyde dehydrogenase (ALDH) activity contribute to neuronal death and AD progression ^36^ . Coupling of rpAD TauO to metabolic enzymes suggests that oligomers in this subtype may directly perturb detoxification and energy metabolism rather than primarily engaging translational or lysosomal machinery. The Rho-GTPase/actin-cytoskeleton signature aligns with reports that tau and tau oligomers remodel actin networks and spine structure, thereby contributing to synaptic failure ^37^. MCODE-based PPI clustering further underscored these patterns. In controls, major modules corresponded to translation and proteostasis, mitochondrial translation, DNA/chromosome organization, intermediate filament architecture and ECM/laminin-integrin networks. spAD modules were dominated by a compact retromer/retrograde transport cluster, with additional coagulation and mRNA-processing subnetworks. rpAD yielded essentially a single, dense metabolic module centered on carboxylic-acid and nucleotide metabolism, while the shared spAD-rpAD subset formed an amino-acid catabolic cluster. These network-level differences support a model in which TauO in rpAD is embedded in a qualitatively distinct cellular context, dominated by metabolic stress and cytoskeletal remodeling.

These alterations support the concept that TauO proteins are not passive aggregates but copurify with functionally relevant proteins, potentially reflecting altered associations that exacerbate cellular stress and degeneration in rpAD ^38^.

Among individual proteins, several interactors showed increased association with TauO in rpAD relative to control and spAD, highlighting a distinct molecular environment underlying the rapid phenotype. SERPINA1 (α1-antitrypsin) a serine protease inhibitor with established neuroinflammatory relevance and known to exhibit disease-specific isoform shifts in AD CSF and brain tissue ^37^. was more enriched in rpAD TauO complexes. Its increased binding suggests heightened engagement of protease-regulatory or extracellular matrix modifying pathways in rpAD. Similarly, ALDH9A1 and related aldehyde dehydrogenases were markedly more abundant in rpAD TauO fractions. Although direct evidence for ALDH9A1 tau interactions is limited, multiple studies implicate ALDH dysfunction in accumulation of toxic aldehydes and neuronal vulnerability in AD ^39,40^. Their increased association with TauO aligns with the strong aldehyde-metabolism signature in rpAD and suggests that TauO in this subtype may sequester or dysregulate detoxifying enzymes at sites of metabolic stress.

Cytoskeletal regulators DPYSL2 (CRMP2) and DPYSL3 (CRMP4) were also elevated in rpAD TauO preparations. CRMP proteins modulate axon guidance, microtubule stability and autophagy, and CRMP2 dysregulation has been implicated in AD pathogenesis ^41^. Their increased recruitment into rpAD TauO suggests enhanced interaction between oligomeric tau and “tau-like” microtubule-associated proteins, potentially amplifying cytoskeletal destabilization in this aggressive clinical variant. NFASC (neurofascin), an axon initial segment and node-of-Ranvier adhesion molecule essential for saltatory conduction ^42^, was similarly enriched, raising the possibility that rpAD TauO may more strongly target axoglial junctions, which could contribute to the rapid functional decline characteristic of rpAD. Another rpAD-specific increase was observed for MAPRE3 (EB3), a microtubule-plus-end tracking protein that stabilizes dynamic microtubules and interacts with tau ^43^. Increased MAPRE3 binding to TauO in rpAD suggests altered regulation of microtubule plus-end dynamics, suggesting a model in which distinct tau oligomer conformations might perturb cytoskeletal remodeling more strongly in rpAD than in spAD.

In contrast, MRPL17, a mitochondrial ribosomal protein, showed reduced association with TauO in rpAD, implying weakened coupling between TauO and mitochondrial translation machinery. Loss of these interactions may reflect impaired mitochondrial resilience or reduced capacity to compensate for metabolic stress in rpAD. Likewise, the terminal complement component C9, implicated in synaptic pruning and neuroinflammation in AD ^44^, was diminished in rpAD TauO complexes, suggesting reduced engagement of complement-related pathways that may otherwise moderate or modulate early immune responses.

Collectively, these quantitative differences show that rpAD TauO is characterized not only by stronger coupling to metabolic and cytoskeletal pathways but also by weakened interactions with mitochondrial and complement components, supporting the view that TauO exists in a qualitatively distinct molecular environment in rpAD. These altered associations may contribute to the metabolic vulnerability, cytoskeletal destabilization and accelerated neurodegeneration that define this clinical subtype. While proteomics was performed on n=3 rpAD samples due to material constraints, the convergence across independent analytical layers (PTMs, TEM, proteotoxicity, network architecture) provides confidence in the robustness of the subtype-specific differences. As a cross-sectional postmortem study, the work provides a snapshot of late-stage TauO biology; future longitudinal and mechanistic studies will be required to determine when these subtype-specific interactome differences arise and how they influence disease progression. This study analyzed a rare rpAD cohort, and the limited availability of well-characterized cases constrained the sample size, particularly for proteomics (n=3). Because immunoprecipitation captures co-purifying proteins, the interactome reflects both direct and compartment-proximal associations. Additionally, the cross-sectional postmortem design provides a late-stage snapshot of TauO biology, and longitudinal analyses will be needed to define when subtype-specific interactome divergence emerges. Despite these constraints, the convergence of PTM, ultrastructural, toxicity, and pathway-level proteomic differences strongly supports the robustness of the subtype distinctions reported here.

## Conclusion

To the best of our knowledge our work provides the first systematic comparison of brain-derived tau oligomers (TauO) from spAD and rpAD . The data consistently demonstrate that rpAD is defined by a structurally and biochemically distinct TauO species. rpAD TauO displayed compact ultrastructural morphology, the highest levels of disease-associated phosphorylation (pS396, pS422), and a fundamentally reorganized interactome. Whereas control and spAD TauO retained robust associations with translation, proteostasis, mitochondrial respiration, and vesicle-trafficking networks, these modules were absent in rpAD. Instead, rpAD TauO showed selective coupling to aldehyde detoxification, amino-acid and carbon metabolism, and cytoskeletal remodeling pathways. Increased recruitment of proteins such as SERPINA1, ALDH9A1, MAPRE3, DPYSL2, DPYSL3, and NFASC alongside depletion of mitochondrial (MRPL17) and complement components (C9) points to intensified metabolic stress and cytoskeletal destabilization in rpAD. These findings support the concept that rpAD harbors a distinct tau oligomer “strain” that engages a different cellular environment and may underlie the fulminant clinical course. These findings provide a molecular framework that advances our understanding of TauO heterogeneity across AD subtypes and may inform future efforts toward subtype-specific biomarkers or therapeutic approaches.

## Declarations

### Ethics approval and consent to participate

All human brain tissues were obtained from the HUB-ICO-IDIBELL Biobank, the Biobank of Hospital Clinic-IDIBAPS (Spain), and the Department of Neuropathology at the University Medical Center Göttingen (Germany). All procedures complied with national legislation and institutional requirements and were approved by the respective ethics committees (Spain: HUB-ICO-IDIBELL Biobank, Hospital Clinic-IDIBAPS; Germany: University Medical Center Göttingen, protocols Nr. 1/11/93 and Nr. 9/6/08).

### Consent for publication

All authors have approved the contents of this manuscript and provided consent for publication.

### Data availability

The datasets generated during the current study are available from the corresponding author on reasonable request

### Author contributions

T.S. conceived and designed the study, performed experiments, analyzed data, and drafted the manuscript. W.M. performed electron microscopy for all samples. M.S. contributed to study design and manuscript writing. A.d.S.C. contributed to data analysis and proofreading. C.T. and C.S. provided and characterized control brain samples. P.H. and S.G. contributed clinical information and cohort characterization for the rpAD cases. S.Z. assisted with data acquisition and analysis. E.R. supported clinical and cohort characterization. C.S. contributed to characterization of control brain samples. O.A. performed mouse experiments. I.F. contributed to AD sample characterization and manuscript preparation. M.H. and T.F.O. provided supervision, guidance, and critical manuscript revision. N.Y. conceived the idea, contributed to study design, data interpretation, and manuscript revision. I.Z. provided overall supervision, secured funding, and critically revised the manuscript. All authors read and approved the final manuscript.

