## Supplemental File for "Tau oligomer heterogeneity and associated protein profile in slowly versus rapidly progressive Alzheimer’s disease"

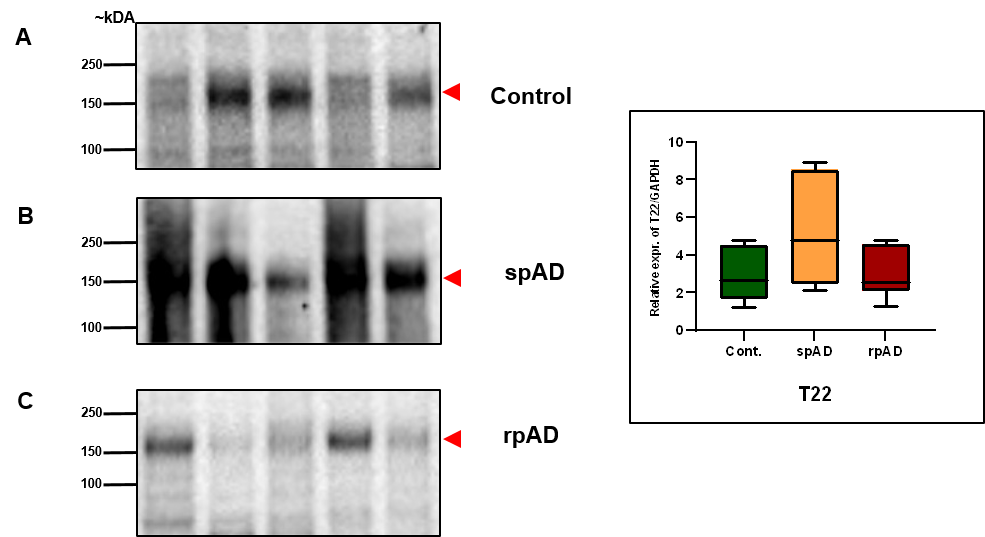

**Figure 1S: Western blot analysis of tau species in PBS crude lysates across the control spAD and rpAD groups.** Representative Western blot images (left) display HMW TauO detected in the control (A), spAD (B), and rpAD (C) groups, with the quantification of band intensities shown in the adjacent bar graph (right). Compared with the control and rpAD groups, the spAD group presented an increase in the intensity of HMW tau species. (n=15 (5 control, 5 spAD, 5 rpAD)).

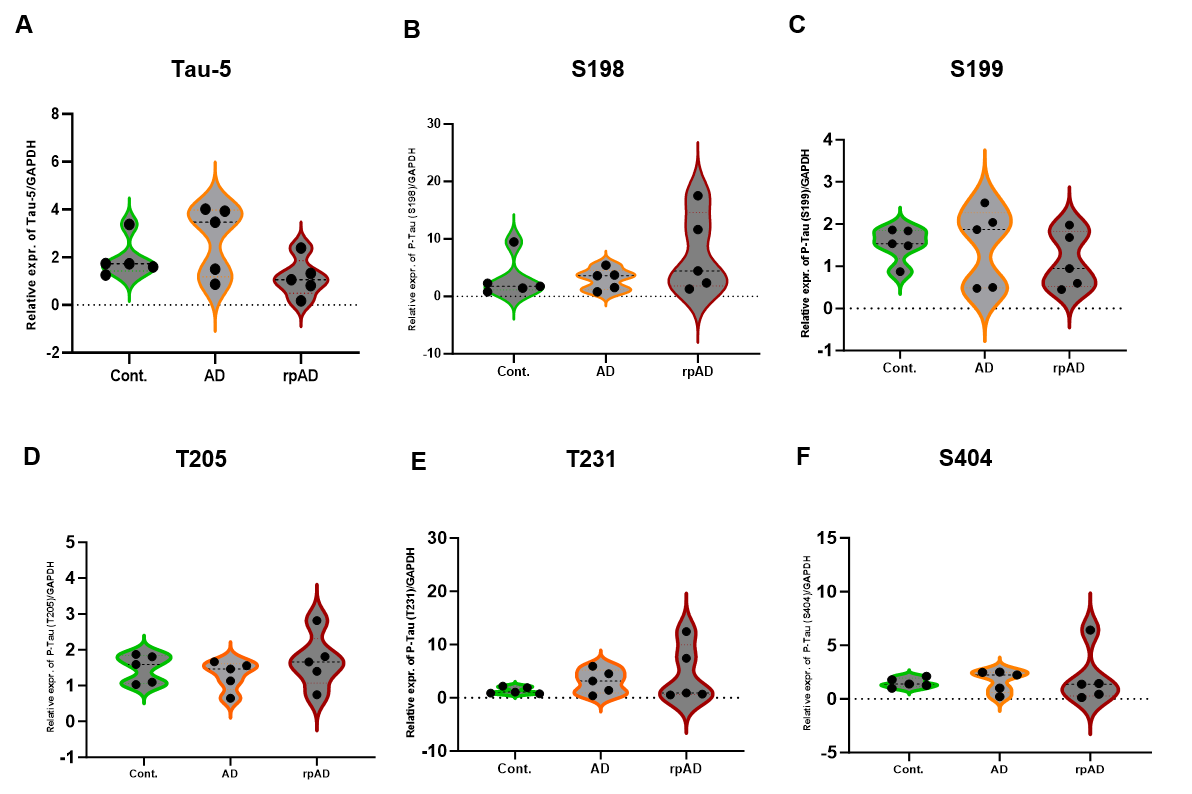

**Figure 2S. Phosphorylation of soluble tau at selected epitopes in AD subtypes and controls.**

Violin plots showing Western blot quantification of total tau (A, Tau-5) and phosphorylation at key residues (B, S198; C, S199; D, T205; E, T231; F, S404) in urea thiourea brain lysates from control, spAD, and rpAD patients. No statistically significant differences were observed across groups. Data represent individual cases quantified relative to total protein loading (n=15 (5 control, 5 spAD, 5 rpAD)).

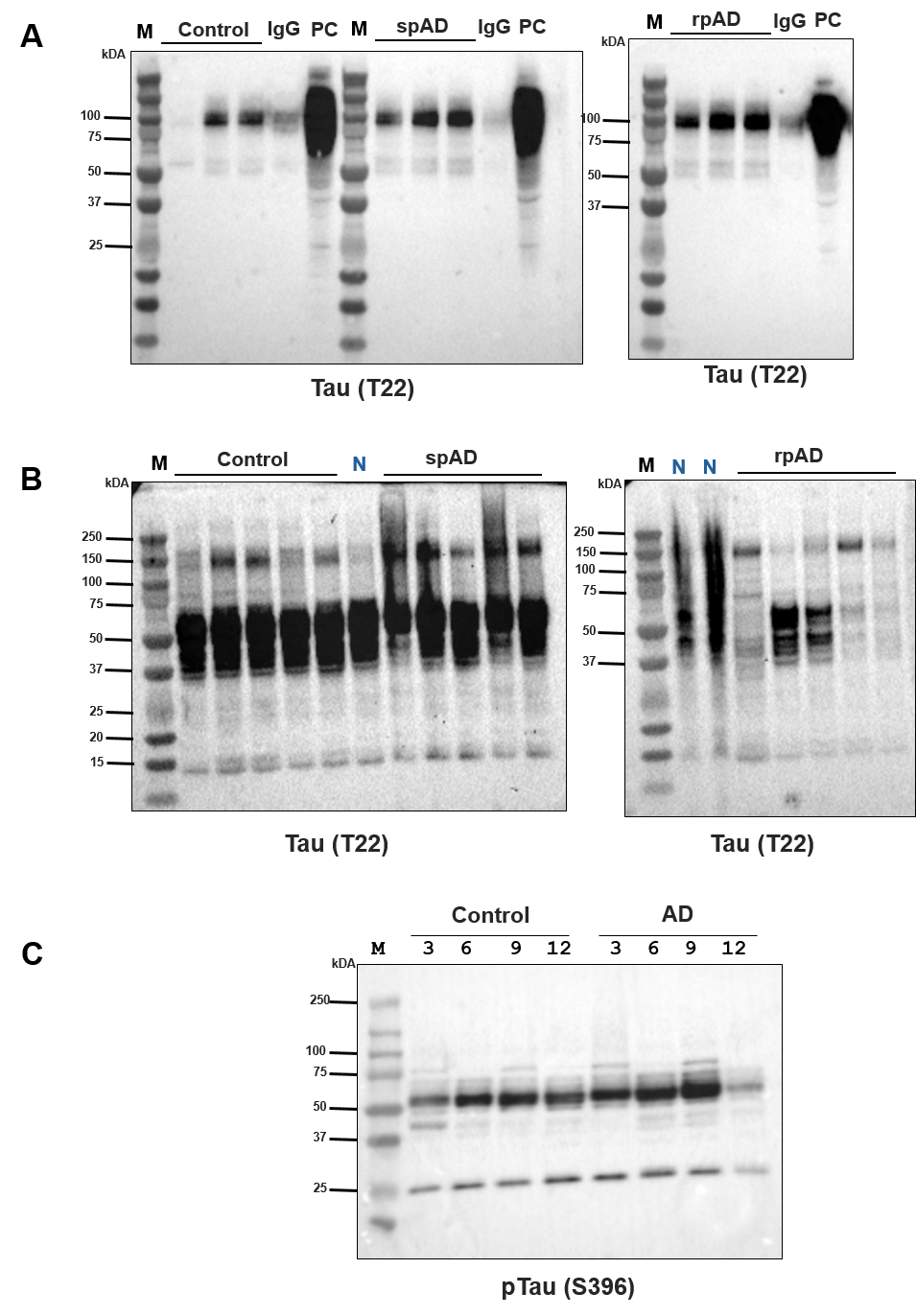

Figure 3S. (A) Uncropped T22 Western blot corresponding to Figure 1 in the main manuscript, showing Tau oligomer immunoprecipitation from control, spAD, and rpAD brain samples, including IgG pull-down controls. (B) Uncropped pTau (T22) Western blot corresponding to Supplementary Figure 1S, showing full-length blot. Lanes labeled “N” represent non-AD samples that were included as processing controls but were not analyzed or interpreted in the manuscript. (C) Uncropped S396 Western blot corresponding to Figure 3D in the main manuscript.

Table 1S: Summary of cases. Control; spAD: sporadic AD; rpAD: rapid Alzheimer disease; N: normal; Braak NFT: Braak neurofibrillary tangle pathology (0-VI); TAP: Thal αB phase (1-5); CERAD: Consortium to Establish a Registry for Alzheimer disease (C0-C3); NIA score: National Institute on Aging (A0-A3); PMI: postmortem delay. ABC categorization: A: TAP: amyloid score; B: Baak NFT pathology; C: CERAD (CERAD-NIA-AA score)

| **No.** | **Case** | **Clinical diagnosis** | **Age** | **Gender** | **Disease duration (M)** | **PMI hours:minutes** | **Braak NFT** | **CERAD-(NIAAA score)** | **Thal** |
| --- | --- | --- | --- | --- | --- | --- | --- | --- | --- |
| 1 | Control 1 | N | 73 | M | - | 91 | - | A0-1B0 |  |
| 2 | Control 2 | N | 62 | M | - | 48 | - | A0-1 B0 |  |
| 3 | Control 3 | N | 59 | M | - | 76 | - | A0-1 B0 |  |
| 4 | Control 4 | N | 62 | M | - | 89 | - | A0-1 B0 |  |
| 5 | Control 5 | N | 61 | M | - | 80 | - | A0-1 B0 |  |
| 6 | spAD 1 | AD; amyloid angiopathy | 81 | F | - | 05:15 | V | C |  |
| 7 | spAD 2 | AD | 82 | F | - | 01:45 | V | B |  |
| 8 | spAD 3 | AD; amyloid angiopathy ;aging related tau astrogliopathy | 75 | M | - | 09 | V | C |  |
| 9 | spAD 4 | AD ;LBD amygdala | 82 | M | - | 3:15 | V | C |  |
| 10 | spAD 5 | AD, LBD, cerebrale amyloid angiopathy | 78 | M | 6 | 24 | V | C |  |
| 11 | rpAD 1 | AD | 84 | M | 3 | 72 | V | B | 3 |
| 12 | rpAD 2 | AD | 72 | F | 2 | 144 | VI | C | 3 |
| 13 | rpAD 3 | AD, cerebrale amyloid angiopathy | 71 | F | 12 | 96 | VI | C | - |
| 14 | rpAD 4 | AD | 77 | M | 12 | 24 | VI | C | 4 |
| 15 | rpAD 5 | AD | 80 | F | 20 | 264 | V | C | - |
